# Design guidelines for α-helical peptides that self-assemble into transmembrane barrel pores killing antibiotic-resistant bacteria

**DOI:** 10.1101/2022.05.09.491086

**Authors:** Rahul Deb, Ivo Kabelka, Jan Přibyl, Edo Vreeker, Giovanni Maglia, Robert Vácha

## Abstract

De novo design of peptides that self-assemble into transmembrane barrel-like nanopore structures is challenging due to the complexity of several competing interactions involving peptides, lipids, water, and ions. Here, we develop a computational approach for the de novo design of α-helical peptides that self-assemble into stable and large transmembrane barrel pores with a central nano-sized functional channel. We address the lack of existing design guidelines for the de novo pore-forming peptides and propose 52 sequence patterns, each of which can be tailored for different applications using the identified role of its residues. Atomic force microscopy, channel electrical recording, leakage of small fluorescent molecule and transport of macromolecule experiments confirm that the designed peptides form stable, large, and functional barrel-shaped nanopores in model membranes. The custom-designed peptides act as potent antimicrobial agents able to kill even antibiotic-resistant ESKAPE bacteria at micromolar concentrations, while exhibiting low toxicity to human cells. Peptides and their assembled nanopore structures can be similarly fine-tuned for other medical and biotechnological applications.

## INTRODUCTION

De novo design of protein sequences that fold and spontaneously assemble into a target structure with the desired function is a crucial test of our current knowledge about protein self-assembly and the ability to develop protein-based tools. Designed protein tools include small peptides, which can modulate the permeability of biological membranes by self-assembling into transmembrane barrel pore (TBP) structures, i.e., transmembrane peptide bundles with a central channel/pore of a few nanometers width, capable of conducting small molecules across the lipid membranes^1,2^ (Figure 1a). TBP-forming peptides have applications in both biotechnology and medicine^3,4^, including antimicrobial peptides (AMPs)^5,6^. Pore-forming AMPs can directly kill bacteria, including multidrug-resistant strains, by rapid permeabilization of bacterial membranes,^7^ and bacteria develop resistance slowly against such AMPs^8^, suggesting that these peptides could become the next-generation antibiotics^9^. However, despite the discovery of thousands of AMPs, alamethicin remains the only known natural AMP which was conclusively shown to form TBP^10^.

**Figure 1.**
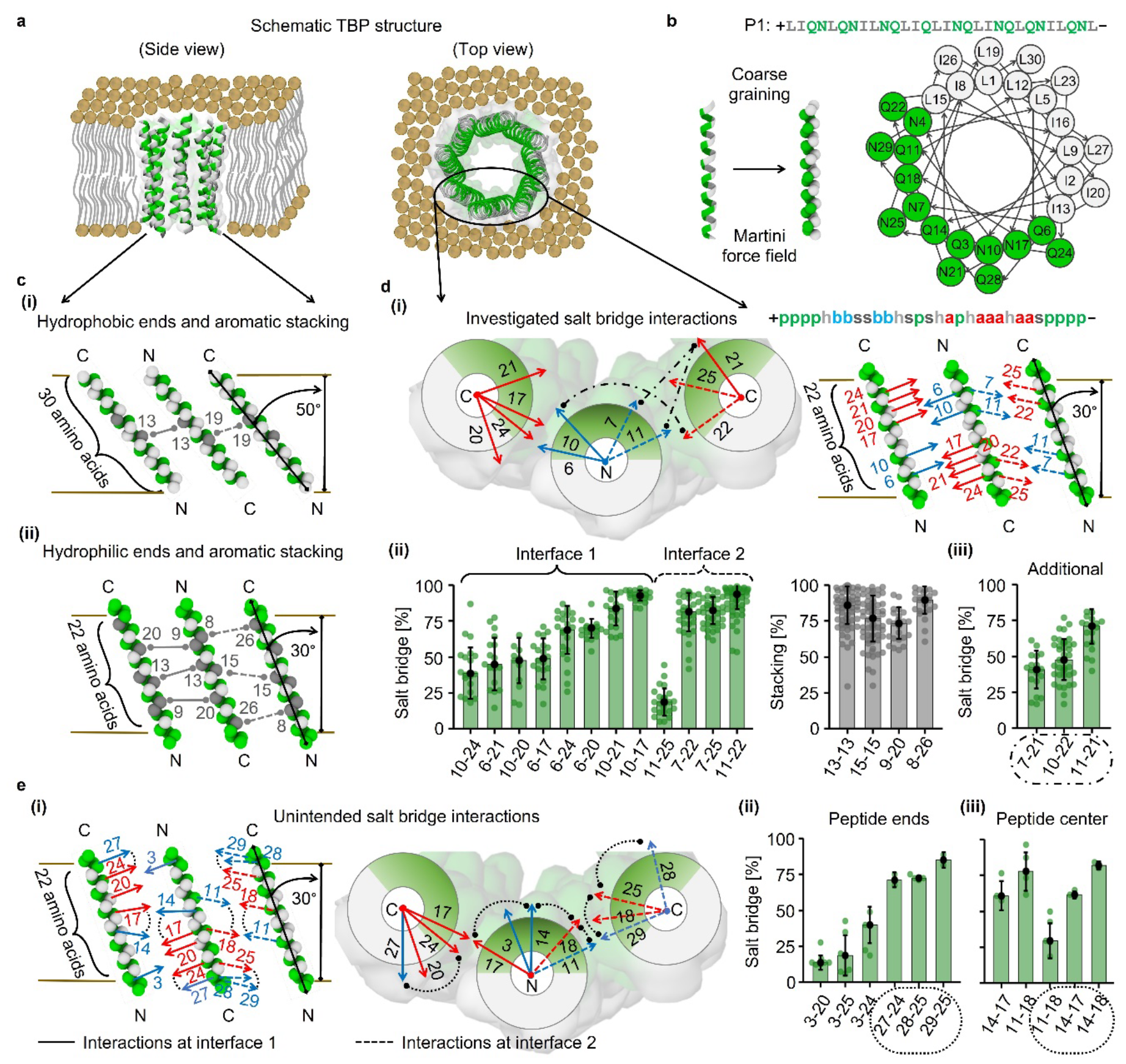
Optimized intermolecular aromatic stacking and salt bridge interactions for stabilization of large TBPs. (a) Schematic representation of an octameric TBP formed by α-helical (ribbons) peptides (30 amino acids). (b) Sequence, helical wheel, and AA-to-CG representations of amphipathic α-helical peptide P1 (Table S1). (c) Schematic illustration of residues forming aromatic stacking interactions. (d)(i) Schematic illustration of the investigated salt bridges and corresponding sequence pattern, composed of polar (p: Q/N), hydrophobic (h: L/I), basic (b: R), acidic (a: D), and aromatic (s: W/F) residues. Basic/acidic and aromatic positions can also be modified with polar and hydrophobic residues, respectively. Stability of intermolecular (ii) salt bridges and aromatic stacking interactions, and (iii) additional salt bridges (dotdash lines) involving two neighboring peptide-peptide interfaces. (e)(i) Schematic illustration and (ii-iii) stability of unintended inter- and intramolecular (dotted lines) salt bridges involving (ii) the charged polar ends of neighboring transmembrane peptides and (iii) peptide center. Basic and acidic residues forming salt bridges are shown with blue and red arrows. Interactions are shown between three antiparallel neighboring transmembrane peptides (tilted by 50°/30° with respect to membrane normal; horizontal lines demarcate membrane leaflets) displaying two representative peptide-peptide interfaces from octameric TBP. In the plots, each point represents one CG simulation showing the percentage of designed/intended interaction contacts averaged over 51 μs simulation (Tables S2 and S3). Color coding: peptide backbone hydrophilic and hydrophobic residues/patches in green and white; aromatic residues in silver; membrane lipid heads in ochre, and tails in gray.

De novo design of TBP-forming peptides is challenging because of the number of competing interactions that affect the stability of pore structure in the lipid membranes.^11^ Peptide-peptide interactions, which are necessary to stabilize TBP structure, compete with peptide-membrane and peptide-water/ion interactions. Due to such a complex environment, the self-assembled pore structures have been shown to be highly sensitive to peptide amino acid composition, with a single-residue mutation capable of destabilizing the pore structure^12^. There are only a few examples of de novo designed helical structures similar to TBPs - e.g., peptides forming narrow ion channels: (LSSLLSL)_3_ peptides^2^, Zn^+2^/H^+1^ antiporter^13^, and coiled-coils initially designed to form water-soluble barrel pores^14^; helix-loop-helix forming double-barrel pores^15^; and redesigned channel-containing natural proteins^16–21^. However, the generalized design principles are still missing, and the role of individual residues in these structures remains either unknown or involved in specific interactions, restricting the introduction of possible mutations for different applications. Experimental determination of residue role is challenging because it requires high spatial and temporal resolution of potentially metastable nanostructures^22^. Computer simulations can provide the required atomic resolution^23,24^.

Here, we perform molecular dynamics (MD) simulations to determine the role of amino acids at each sequence position for the stabilization of large TBPs. Based on simulations, we formulate a set of design guidelines and propose four groups of sequence patterns for α-helical peptides that are able to form stable TBPs with a central water channel. We experimentally verify peptide TBP formation by a combination of channel electrical recordings, fluorescence, and atomic force microscopy (AFM) experiments. Provided guidelines enable custom design of peptide nanopores tunable for specific functions. We use the obtained knowledge to de novo design pore-forming potent AMPs that are capable of selectively killing pathogenic resistant Gram-positive and Gram-negative bacteria while sparing human cells, suggesting their potential antibiotic functions, and implying that peptides can be similarly tuned for other medical and nanobiotechnological applications.

## RESULTS

### Computer simulations

To analyze the function of each residue in TBP stabilization/destabilization, we designed ∼150 model peptides with systematically modified sequences, and tested the ability of each peptide to stabilize TBP in MD simulations using the coarse-grained (CG) Martini 2.2 force field^25^ (see Methods in SI for details). In short, we started with a structure consisting of a preformed membrane pore lined by 8 transmembrane peptides, from which the system could rearrange in an unbiased manner. Depending on the strength of specific intermolecular peptide-peptide interactions, we observed either pore closure, deformation, or stabilization of TBPs. We assumed that once the TBP structure becomes the lowest energy state, peptides would be able to spontaneously assemble into such TBP structures after crossing the kinetic barriers^26^. We chose to design single-ring octameric TBPs (i.e., composed of 8 peptides), as these are the smallest nanopore structures that allow the passage of organic molecules bigger than water/ions^15,16,18^. Note, however, that the designed peptides can form larger TBPs if the peptide-peptide interactions are able to adapt such arrangements. We focused on amphipathic α-helical peptides composed of 30 amino acids, an approximate helix length required to span the thickness of a common model membrane made of 1-palmitoyl-2-oleoylsn-glycero-3-phosphocholine (POPC) lipids. We selected antiparallel peptide arrangement in TBPs, which is favored over the parallel arrangement in helix self-assembly and pore formation^23,27,28^. The pore was considered ‘stable’, ‘deformed’ or ‘closed’ if the number of transmembrane peptides (in the pore) at the end of a 51 μs long unbiased simulation run was 8, 7-4 or < 4, respectively.

### Hydrophobic residues

Close packing of hydrophobic side chains (i.e., knobs-into-holes interactions^2,14^) and aromatic residues support oligomerization of transmembrane helices^29,30^. To investigate the pore-stabilizing effects of these interactions, we studied 6 peptides P1-P6 (Table S1 and Figure S1a), in which hydrophobic residues were systematically varied. We simply started with a 50% amphipathic sequence, composed of highly hydrophobic and hydrophilic residues with a strong propensity for α-helix formation. Peptide hydrophobic patch was made of leucine (L) and isoleucine (I) to maintain strong interactions with the membrane, and the hydrophilic patch was made of glutamine (Q) and asparagine (N) to maintain high polarity in the pore lumen (Figure 1b). Regardless of the changes in the hydrophobic content, P1-P3 did not stabilize the pore (P2 and P3 were 23% and 77% hydrophobic). Substituting L/I to aromatic tryptophan (W) and phenylalanine (F) in P4 and P5 improved pore stability. An expanded W-patch in P6 (L/I to W in P3) resulted in stable TBP due to increased peptide-peptide interactions, although the pore cavity was too narrow for a continuous water channel (Figure S1b).

We next investigated specific distributions of aromatic residues in the middle of the sequence using peptides P7-P15 (Table S1). We found that these residues need to be in specific positions to promote the formation of intermolecular aromatic π-π stacking interactions to enhance TBP stability. Aromatic residues that were placed close to each other on a helical sequence led to intramolecular interactions (within the same helix) that interfered with the designed intermolecular interactions (between neighboring helices) and did not stabilize TBPs (Figure S1c). Optimal positions for residues forming intermolecular interactions depend on transmembrane peptide tilt in the pore (i.e., the angle between peptide helical axis and membrane normal), which is in turn determined by the hydrophobic mismatch between membrane hydrophobic core and peptide hydrophobic patch^23,27^. With a hydrophobic patch spanning over the whole length of the peptides (i.e., 30 residues), P14 and P15 adopted a tilt of 50° and the optimal positions were 13 and 19 for single stacking interaction on each peptide-peptide interface in TBP (Figures 1c(i) and S1d). However, these interactions were not sufficient to stabilize TBPs. While stronger W-stacking resulted in a deformed pore after 51 μs with P14, weaker F-stacking did not stabilize the pore after 33 μs with P15 (Figures S1e-f). An additional W residues at position 16 (in P10) could hinder 13-13 and 19-19 intermolecular stacking interactions by forming intra-molecular ones, eventually closing P10 pore after 24 μs (Figure S1c). Thus, the intricate interplay of aromatic residues in inter- and intramolecular interactions requires their careful positioning in the sequence for TBP stabilization.

Peptide hydrophilic ends can improve TBP stability by an enhanced preference for the transmembrane orientation due to electrostatic interactions and hydrogen bonding with lipid headgroups. Moreover, such interactions can be utilized for lipid specificity, e.g. cationic ends can increase peptide selectivity for the negatively charged bacterial membranes over zwitterionic human membranes^5,31,32^. Therefore, we shortened the hydrophobic patch from 30 to 22 residues (in the middle of the sequence), leaving 4 hydrophilic residues at each end to interact with lipid headgroups (Table S1). We tested two types of cationic ends using arginine (R) residues (i.e., NQRR- and RRQN-) and found that the peptides P16 and P17 with R-ends resulted in deformed pores, while the parent peptide P1 did not stabilize the pore at all. C-terminus amidation (-NH2) decreased pore stability; P17-NH2 closed the pore and P16-NH2 produced a deformed pore. These results suggest that cationic ends can stabilize transmembrane orientation of peptides in TBP, and interactions between the oppositely charged neighboring peptide termini (i.e., N-terminus NH3^+^ and C-terminus COO^-^) promote pore stability, as reported previously^23,33^.

We next investigated optimal aromatic stacking interactions to improve TBP stability in presence of hydrophilic and cationic residues at the peptide ends using peptides P18-P27 (Table S1). With a reduced hydrophobic patch spanning 22 residues, peptides adopted a decreased tilt of 30° in TBP and the optimal positions were 13 and 15 for single stacking and 8, 9, 20, and 26 for double stacking interactions on each peptide-peptide interface in TBP (Figures 1c(ii) and S1g). P20-P21 with cationic ends and 13-13 plus 15-15 single stacking resulted in deformed TBPs, whereas P18-P19 lacking hydrophilic ends closed the pore. Similarly, P24-P27 with cationic ends and 9-20 plus 8-26 double stacking interactions stabilized TBPs, whereas P22-P23 lacking hydrophilic ends closed the pore. As expected, the pore-stabilizing effect of double stacking interactions per interface was stronger than single ones.

### Hydrophilic residues

Apart from the (water-mediated) hydrogen bonds between peptides, salt bridge interactions can contribute to pore stability^34^. We analyzed TBP stabilization with salt bridges between R and aspartic acid (D), designed at all possible hydrophilic positions using 75 model peptides based on templates P20-P21 and P24-P25 (with single and double stacking interactions per peptide-peptide interface) (Tables S2 and S3). Peptides were designed with either 2 (groups A and B) or 4 (groups C and D) salt bridges on each peptide-peptide interface in TBP (peptide nomenclature in supplementary texts). Charged residues at positions 6, 10, 17, 20, 21, and 24 were tested on the first interface, and at 7, 11, 22, and 25 on the second interface (Figure 1d(i)). Overall, pore stability increased in presence of salt bridges, and the stability of tested salt bridge residue pairs increased as follows: 10-24 < 6-21 < 10-20 < 6-17 < 6-24 < 6-20 < 10-21 < 10-17 on the first interface, and 11-25 < 7-22 < 7-25 < 11-22 on the second interface (Figure 1d(ii)).

Other than the designed/intended intermolecular interactions, we also observed the formation of unintended intermolecular salt bridges in the pore: 7-21, 10-22, and 11-21 involving the neighboring interfaces (Figure 1d(iii)) and 3-20, 3-24, and 3-25 involving the neighboring peptide ends (Figures 1e(i-ii)), which increased TBP stability. Similarly, we identified the formation of unintended intramolecular salt bridges 14-17 and 11-18 at the peptide center, and 27-24, 28-25, and 29-25 at the peptide C-terminus end (Figure 1e, (i) and (iii)) that decreased the stability of designed intermolecular salt bridges (in particular 6-24 and 7-25).

Extended ladder of salt bridges, known as electrostatic “charge zipper”, was reported to promote the self-assembly of transmembrane helices^34^. In our design, 4 intermolecular salt bridges on each peptide-peptide interface (groups C and D) acted as such an intermolecular charge zipper (Figure 1d(i)). Stabilized TBPs made of peptides with charge zippers were more compact and rigid than the TBPs made of 2 salt bridges per interface (i.e., groups A-B peptides without charge zipper) in three types of mutations investigated using 33 peptides.

The first type of mutations was R to lysine (K) substitutions and/or increased peptide net charge by adding cationic residues at the polar ends and/or on the polar faces (Figure 2a and Table S4). With these we tested the possible increase in bacterial selectivity because K residues interact stronger with anionic phospholipids (rich in bacterial membranes) than with the zwitterionic phospholipids (rich in human membranes)^5,31,32,35^. Stability of TBPs and salt bridges remained intact after K-substitutions, irrespective of the number of interactions per interface. However, cationic residues at the polar ends of groups C and D peptides resulted in the formation of unintended inter- and intramolecular salt bridges (Figures 1e(i-ii)), which caused subtle changes in the stability of designed salt bridges. For group A peptides, unintended intermolecular salt bridges formed after adding cationic residues at the polar ends and/or faces caused substantial deformations of the designed interactions (Table S5). Although the overall pore stability remained unaltered, these results suggest that charged residues should be included cautiously.

**Figure 2.**
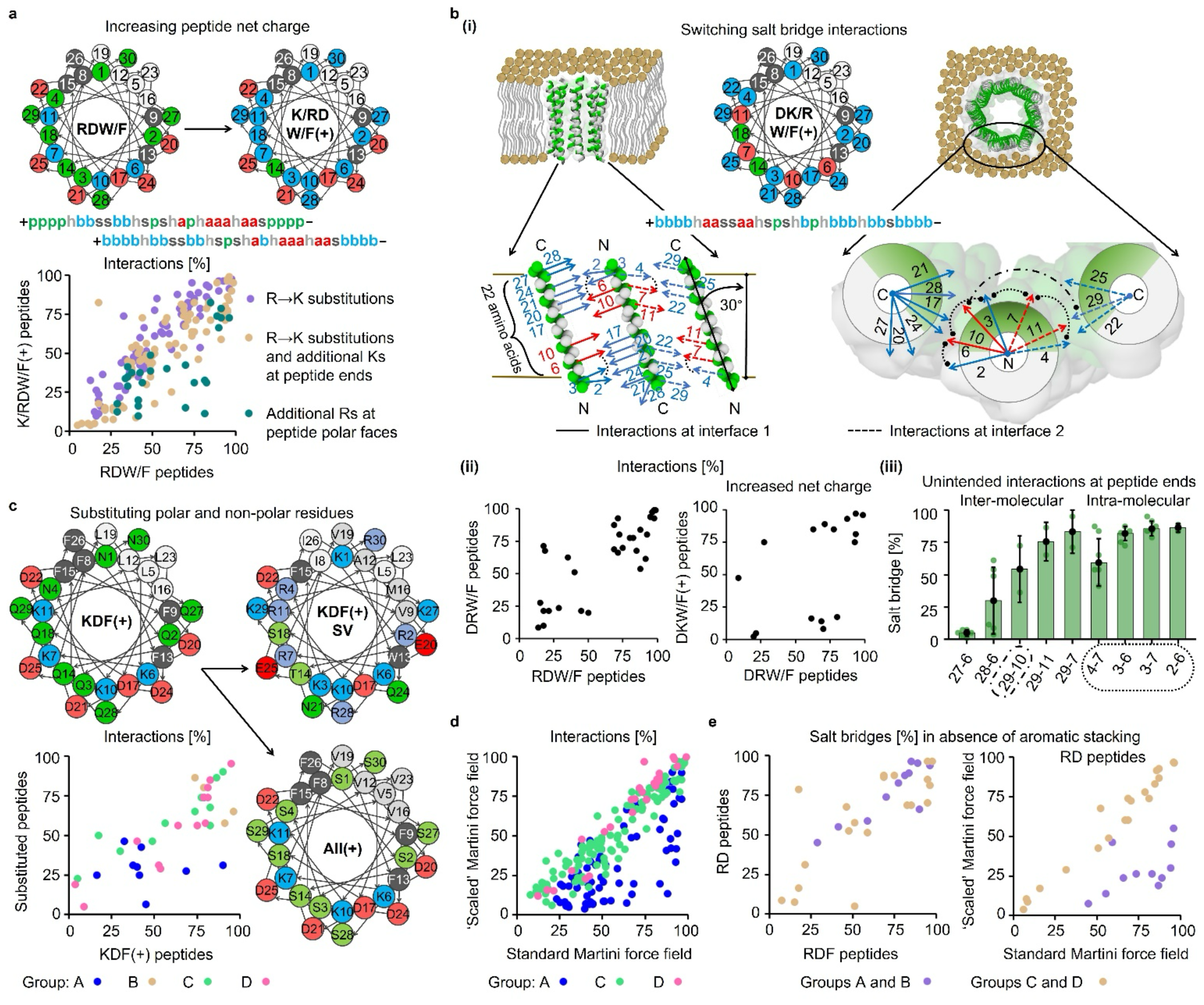
Intermolecular electrostatic charge zippers stabilized octameric TBPs with various mutations. (a) Peptide helical wheels and sequence patterns showing the investigated positions of additional cationic residues in the sequence. Correlation between interactions in TBPs stabilized by RDW/F peptides and their K/R-variants with/without increased net charge (Tables S4 and S5). (b)(i) Peptide helical wheel with sequence pattern, and schematic illustration of the investigated salt bridges between charged residues at switched positions. Sequence patterns are composed of polar (p: Q/N), hydrophobic (h: L/I), basic (b: R/K), acidic (a: D), and aromatic (s: W/F) residues. Basic/acidic and aromatic positions can also be modified with polar and hydrophobic residues, respectively. Salt bridges (basic and acidic residues in arrows) are shown between three antiparallel neighboring transmembrane peptides (tilted by 30° with respect to membrane normal; horizontal lines demarcate membrane leaflets) displaying two representative peptide-peptide interfaces from octameric TBP. (ii) Correlation plots of the stability of individual interactions in TBPs stabilized by RDW/F peptides and their switched salt bridge variants (DRW/F), including increased net charge. (iii) Stability of unintended inter- (including 29-10 in dotdash lines involving two neighboring interfaces) and intramolecular (dotted lines) salt bridges involving the charged polar ends of neighboring transmembrane peptides (Table S6). (c) Peptide helical wheels showing the substituted positions of polar and non-polar residues, and correlation between interactions in TBPs stabilized by KDF peptides and their substituted variants (Table S7). (d-e) Correlation between interactions in TBPs simulated with standard and ‘scaled’ Martini force fields for (d) RDW/F peptides (Tables S2 and S8), and (e) RD peptides without aromatic stacking interactions (Table S9). In the plots, each point represents the percentage of designed/intended interaction contacts averaged over 51 μs simulation. Color coding: peptide backbone hydrophilic and hydrophobic residues/patches in green and white; basic and acidic residues in blue and red; aromatic residues in silver; membrane lipid heads in ochre, and tails in gray.

The second type of mutations was switching the positions of two oppositely charged residues forming a salt bridge pair, i.e., D-R instead of R-D salt bridges (Figure 2b(i)). Testing 2 group-C peptides, we found that switched salt bridges stabilized TBPs equally as the template peptides (Figure 2b(ii) and Table S6). Additional Ks at the polar ends (and R-to-K substitutions) resulted in unintended inter- (27-6, 28-6, 29-7, 29-10, and 29-11) and intramolecular (2-6, 3-6, 3-7, and 4-7) salt bridges (Figure 2b(iii)). TBPs remained stable, but these unintended interactions influenced the stability of a few designed salt bridges (in particular the ones involving positions 7 and 11; Table S6).

The third type of mutation was substitution of polar (Q/N) and non-polar (L/I) residues to serine (S) and valine (V), respectively, to test the influence of the initial LIQN character. We simulated at least one peptide from groups A-D (Figure 2c and Table S7). Compared to the parent peptides, the stability of TBPs and interactions remained largely intact for group C-D peptides, but the pore cavities became narrower due to tight-packing of polar faces (Figures S2d-e). Tight-packing is likely caused by hydrogen bonds between S residues, promoting helix association^36^. SV-substituted peptides from group A-B had reduced interactions and even narrower or even closed pores (Figures S2a-c). Finally, we tested one peptide from group-C with a combination of different amino acids: threonine (T), Q, S, N, alanine (A), methionine (M), V, L, I, W, F, R, K, D, and glutamic acid (E) (Table S7). This All_C16+8 peptide stabilized TBPs for the full length of simulation (Figure S2f), suggesting an enormous space of possible mutations.

### Standard versus ‘scaled’ Martini

The obtained stability of TBPs could be overestimated in simulations with standard Martini force field due to an overestimated protein-protein interactions^37^. Therefore, we verified our results using a ‘scaled’ Martini force field with scaled-down intermolecular pair potential (see Methods for details). TBP stability decreased significantly for peptides from groups A-B, apart from those designed with the most stable 6-20, 7-25, 10-17, and 11-22 salt bridges (Figure 2d and Table S8). However, none of them stabilized TBP without aromatic stacking (W/F-to-L/I substitutions), compared to simulations with standard Martini (Figures 2e and S2g-j, and Table S9). In contrast, TBP stability remained intact for peptides from groups C-D (with charge zippers) even without stacking interactions (Figures 2e and S2k-m). These results confirmed the most stable salt bridges and suggested that group C-D peptides do not require aromatic stacking interactions for TBP stabilization.

### Simulated structure of TBP

The CG simulated structure of our de novo designed TBPs (see Figure 3a for a representative structure of TBP made of RDFA2 peptides) is reminiscent of the alamethicin and (LSSLLSL)_3_ peptide ion channels^1,2^. However, our designed octameric TBPs have a larger tilt angle, antiparallel peptide arrangements, and contain charged and aromatic residues with rationally designed interactions. We verified the local stability of octameric structures and the pore-stabilizing interactions by performing all-atom (AA) simulations using 8 different peptide sequences that stabilized the TBP in CG simulations (Table S10). Starting from (1) CG-equilibrated system and (2) configuration after 51 μs (back-mapped to AA), all the tested peptides maintained stable TBP in 2 μs simulation. The representative AA TBP structures made of RDFA2 and its K-variant KDFA2i+9-NH2 (with increased net charge) show the formation of the designed 6/10-20 plus 7/11-25 R/K-D salt bridges and 13-13 plus 15-15 F-stacking interactions between the peptides (Figure 3b-c). The stability of the designed interactions remained largely unchanged compared to the CG simulations, with very few outliers (Figures S3a-i). As expected, peptide secondary structure remained α-helical for the ∼22 residue central amphipathic patch, while the hydrophilic cationic ends partially unfolded outside the membrane hydrophobic core. The TBPs had a central continuous water-ion channel throughout the 2 μs simulation run (Figures 3d-e and S3j-o). The pore diameter ranged from 10.6 to 14.2 Å, while the minimum pore constriction ranged from 3.6 to 10.2 Å, as calculated by HOLE^38^. Based on AA and CG simulations, the increasing strength of peptide-peptide interactions (per interface) improved the pore stability in the order of groups A < B < C < D. These interactions, however, also led to progressively narrower pore cavities and water channels, as shown in Figure 3f.

**Figure 3.**
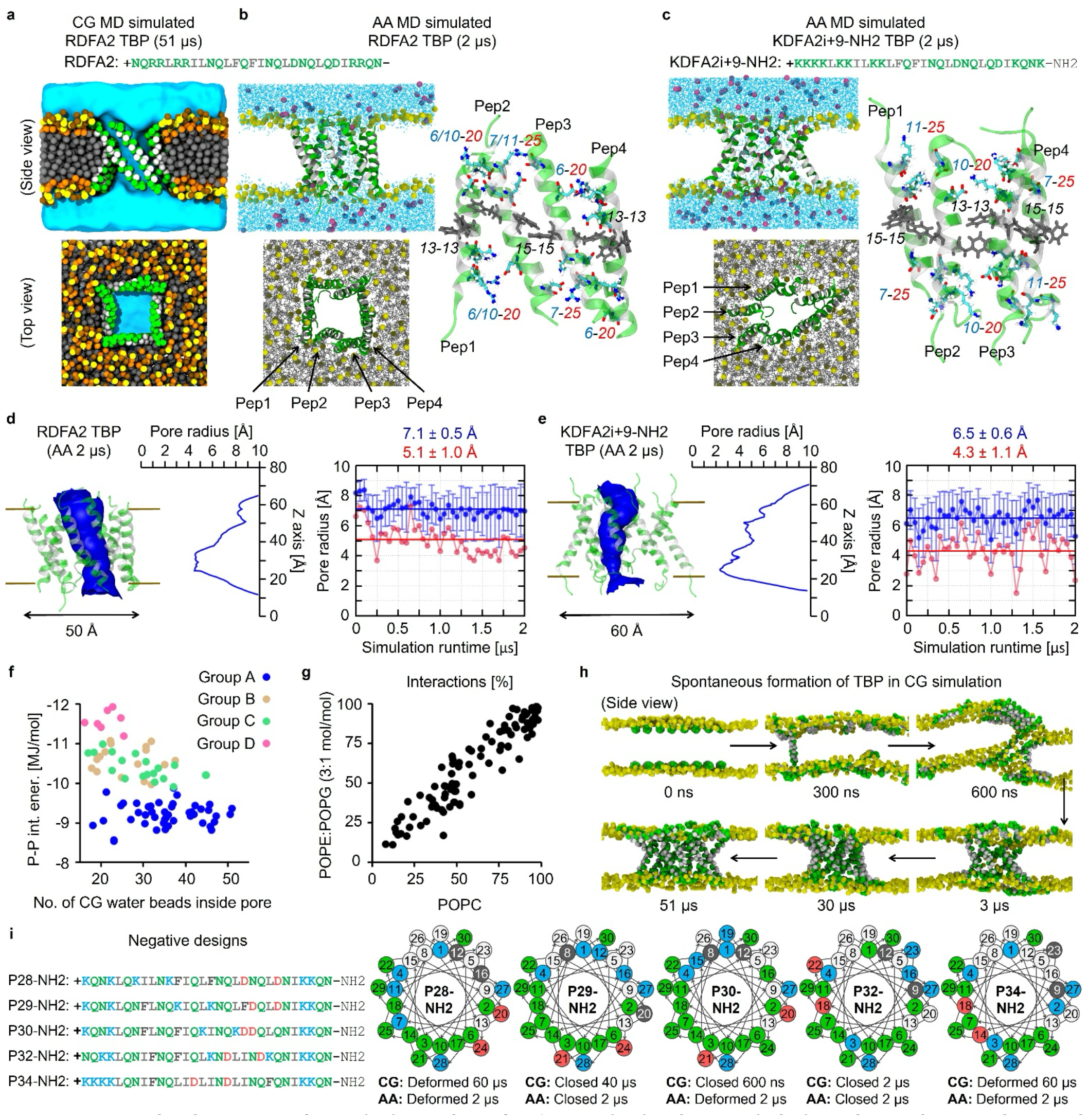
MD simulated structures of TBPs. (a-c) Snapshots after 51 μs CG (in a) and 2 μs AA (in b-c) simulations depicting the central water channels (cyan) through octameric TBPs stabilized by RDFA2 (in a and b) and KDFA2i+9-NH2 (in c) peptides in POPC membrane (Tables S2 and S10). AA simulations were performed by extending back-mapped 51 μs CG simulation. Membrane is represented by only lipid phosphates for the side view clarity. Sodium and chloride ions are shown as blue and pink spheres. Additionally, the designed R/K-D salt bridges and aromatic F-stacking interactions stabilizing the TBPs are shown for four neighboring transmembrane peptides. (d-e) (Left:) The permeation paths in the final 2 μs AA snapshots are shown as blue surface and (Middle:) the radius of the pores along the permeation paths. (Right:) The time-averaged pore radius (in blue) and the minimum pore radius (in red) along the permeation paths over 2 μs simulation runs at 50 ns intervals. Calculations were performed using HOLE^38^. (f) Correlation between the time-averaged numbers of CG water beads inside TBPs and peptide-peptide interaction energies. (g) Correlation between interactions in TBPs simulated in POPC and POPE:POPG (3:1 mol/mol) membranes. Each point represents the percentage of designed/intended interaction contacts averaged over 51 μs simulation. (h) Snapshots from CG simulations showing spontaneous formation of TBP by RDFA2 peptides in POPC:POPG (1:1 mol/mol) membrane (shown with only lipid phosphates for clarity) (Table S11). (i) Sequences and helical wheels of the negative control peptides, comparing TBP stability from CG and AA simulations (back-mapped from CG-equilibrated configurations) in POPC membrane (Tables S12 and S13). Color coding: (a-e h, i) peptide backbone hydrophilic and hydrophobic residues/patches in green and white; (a-c, h) membrane lipid cholines, phosphates, glycerols and tails in ochre, yellow, orange and gray; (i) basic and acidic residues in blue and red; aromatic residues in silver.

### Effect of membrane composition

The stability of TBPs and interactions were very similar in the simulations of zwitterionic POPC bilayer and a charged bilayer made of 1-palmitoyl-2-oleoyl-sn-glycero-3-phosphoethanolamine (POPE) and 1-palmitoyl-2-oleoyl-sn-glycero-3-phospho-1′-rac-glycerol (POPG) lipids (3:1 mol/mol), simple mimics of eukaryotic and bacterial membranes, respectively (Figure 3g). However, the cationic peptide ends had stronger interactions with the phosphates of negatively charged POPG lipids compared to zwitterionic POPC and POPE lipids (Figure S4a). These stronger interactions suggested membrane preference, which was verified by free energy calculations. Peptides had higher affinity to POPE:POPG membrane (∼20 kJ/mol) compared to the neutral POPC membrane (Figure S4b), indicating stronger affinity towards anionic bacterial membranes.

### Spontaneous formation of TBP

Using the designed pore-stabilizing peptides, the following TBP self-assembly process was obtained from the initial membrane surface-adsorbed states: (i) insertion of one peptide into transmembrane state; (ii) insertion of other peptide(s) along the first one, associated with the formation of membrane defects; (iii) formation of a water channel across the lipid membrane; (iv) oligomerization of transmembrane peptides into a disordered (toroidal) pore; and (v) system relaxation into a regular TBP structure with a large internal cavity and continuous water channel (Figure 3h). The self-assembled TBP increased in size over time, becoming even larger than octamers as more peptides were accommodated, suggesting that such large pores may be also stable. The self-assembly process is similar to the reported pore self-assembly processes of AMPs^23,39^. However, spontaneous TBP formation was found to be stochastic already in the limited time scales that we could simulate (Table S11). Such stochastic behavior was anticipated because of the high energy barrier associated with membrane defect/pore formation in Martini force field^40^. Therefore, we also studied systems with facilitated pore formation by initially placing peptides near the membrane hydrophobic core (i.e., at lipid glycerol region) or by an applied membrane thinning potential (Figure S5).

### Negative designs

Before converging to a set of design guidelines for TBP-stabilizing peptides, we verified our findings with negative control peptides that were designed/anticipated not to stabilize TBPs (Figure 3i and Table S12). As a template, we used KDFA2 (K-variant of RDFA2) that stabilized TBP with highly stable 6-20 plus 7-25 salt bridges and 13-13 plus 15-15 aromatic stacking interactions at the first and second peptide-peptide interface in TBP (Table S4). Negative control peptides P28-P34 were designed with similar amphipathicity and amino acid compositions, including K, D, and F, but at positions not expected to form TBP-stabilizing interactions. Additionally, we tested P35 and P36 (Ks) based on peptides P16 and P17 (Rs), respectively, with only cationic residues (i.e., no D and F) at the ends. As expected, all the negative control peptides resulted in either closed or deformed pores, with the only exception of P28. Pore stabilization by P28 is likely a combination of: (i) negatively charged C-terminus interacting with positively charged N-terminus or K-ends of neighboring peptides, (ii) K-ends interacting with lipid phosphates, and (iii) close packing of peptide amphipathic patches, e.g., knobs-into-holes interactions^2,14^. C-terminus amidation and/or L/I-to-A substitutions (which hampered the close packing interactions) reduced the pore stability. Thus P28-NH2 and P28A-NH2 resulted in deformed and closed pores. We verified the above CG results by performing 2 μs long AA simulations, starting with back-mapped CG-equilibrated structures (Table S13). All the tested 9 negative control peptides with amidated C-terminus (including both L/I and A-variants) resulted in either deformed or closed pores, consistent with CG simulations (Figure 3i). The results of negative control peptides together with the pore-stabilizing peptides confirmed that designed peptides stabilize TBPs by a combination of specific intermolecular interactions, including salt bridges, aromatic stackings, close packing of residues, and interactions involving peptide termini/ends.

### Design guidelines

Based on the above *in silico* findings, we developed a set of guidelines for the rational design of α-helical peptides that are able to self-assemble into stable TBPs with a large water channel and an antiparallel peptide arrangement. The guidelines generated 52 sequence patterns with the identified role of each residue (Table S14). Sequence patterns are summarized in 8 templates organized in four groups A, B, C and D, based on the number of interactions between the neighboring peptides (Figure 4). The design guidelines are as follows:

**Figure 4.**
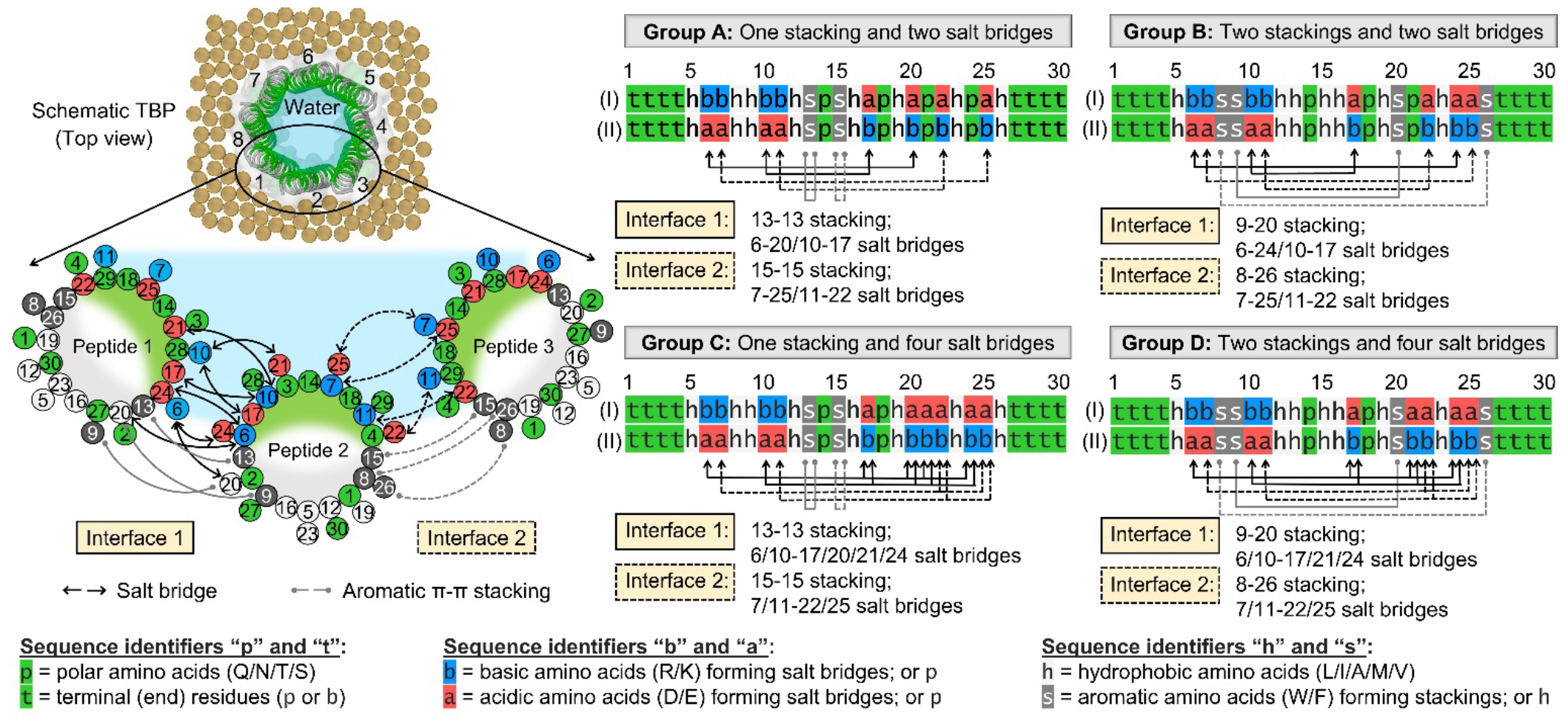
Design guidelines for α-helical peptides that stabilize large TBP structures. Eight sequence templates of length 30 amino acids (with the central hydrophobic patches spanning ∼22 residues) are categorized into four groups (A, B, C, D) based on the total number of aromatic π-π stacking and salt bridge interactions designed on each peptide-peptide interface in TBP. (Left) Schematic illustration of the proposed interactions is shown between three antiparallel neighboring transmembrane peptides (helical wheels) with two representative interfaces. (Right) Sequence templates are defined with identifiers - p, h, b, a, s, and t, characterizing specific amino acids. Two sequence templates (I and II) are proposed for each group with switched positions of charged residues forming salt bridge pairs. Color coding: peptide hydrophilic and hydrophobic residues/patches in green and white; basic and acidic residues in blue and red; aromatic residues in silver; membrane lipid heads in ochre; and water in blue.

1. Proposed TBP-forming peptides are made of 30 amino acids, with a helical amphipathic character spanning across ∼22 amino acids in the middle of the sequence, i.e., a continuous hydrophobic patch in α-helical conformation.
2. Non-polar amino acids forming the hydrophobic patch include A, M, V, L, and I. Polar amino acids forming the hydrophilic patch include T, Q, S, and N. Peptides made of amino acids with high hydrophobicity (L and I) and polarity (Q and N) are preferred for pore stability.
3. Oppositely charged amino acids (cationic R and K, and anionic D and E) within the hydrophilic patch are designed to form intermolecular salt bridge interactions. Either two (groups A and B) or four salt bridges (groups C and D) can be designed on each peptide-peptide interface. TBPs designed with two salt bridges per interface require further stabilization via interactions involving aromatic residues or charged peptide termini/ending residues. Pairs of positions at which the oppositely charged residues form salt bridges are as follows - Group A: 6-20 or 10-17 on the first interface, and 7-25 or 11-22 on the second interface; Group B: 6-24 or 10-17 on the first interface, and 7-25 or 11-22 on the second interface; Groups C and D: the first salt bridge pair chosen from 6-17, 6-20, 6-21, or 6-24, and the second salt bridge pair chosen from 10-17, 10-20, 10-21 or 10-24 for the first interface, and the first salt bridge pair chosen from 7-22 or 7-25, and the second salt bridge pair chosen from 11-22 or 11-25 for the second interface. Each residue should be involved only in one salt bridge interaction. Overall, the stability of salt bridge pairs increases in the following order: 10-24 < 10-20 ≈ 6-21 < 6-17 < 6-24 < 6-20 < 10-21 < 10-17 on the first interface, and 11-25 < 7-22 < 7-25 < 11-22 on the second interface.
4. Aromatic non-polar amino acids (W and F) within the hydrophobic patch are designed to form intermolecular aromatic π-π stacking interactions. Optimal positions are 13 and 15 for a single stacking on each peptide-peptide interface (groups A and C), and 8, 9, 20, and 26 for double stackings on each interface (groups B and D).
5. Four residues at each peptide end are kept polar and can be modified for different applications. Cationic residues at the ends help to stabilize the transmembrane orientation of peptides in TBP. However, for the peptides designed with two salt bridges per interface (groups A and B), the inclusion of such charged residues can alter the stability of designed interactions due to the formation of unintended inter- and/or intramolecular salt bridges. Therefore, the oppositely charged residues should not be combined at the following positions: 6-2/3, 7-3/4, 24-27, and 25-28/29. Furthermore, oppositely charged peptide termini (i.e., N-terminus NH3+ and C-terminus COO-) aid TBP stabilization via attractive interactions between the neighboring antiparallel transmembrane peptides. Note that peptide length can be adjusted for specific functions by extending or shortening both the polar ends.

### Experimental validation of pore formation

To validate the simulation-based design guidelines, we *in vitro* experimentally tested 26 designed peptides for pore-forming activity (Table 1 and Supplementary Texts). We tested at least one pore-stabilizing peptide from each of the four designed groups A to D, including both R and K variant peptides with different net charges (22 peptides in total). In addition, we tested 4 negative control peptides that were designed without the proposed pore-stabilizing interactions. All of these peptides were designed based on simulations, in which only the negative control peptides failed to stabilize TBPs, in contrast to the other peptides. While the peptide N-terminus was always positively charged, we tested both negatively charged and amidated (-NH2) C-terminus. Nineteen of the 26 peptides were soluble in phosphate-buffered saline (PBS), including the 4 negative controls and at least one pore-stabilizing peptide from each group.

**Table 1.**
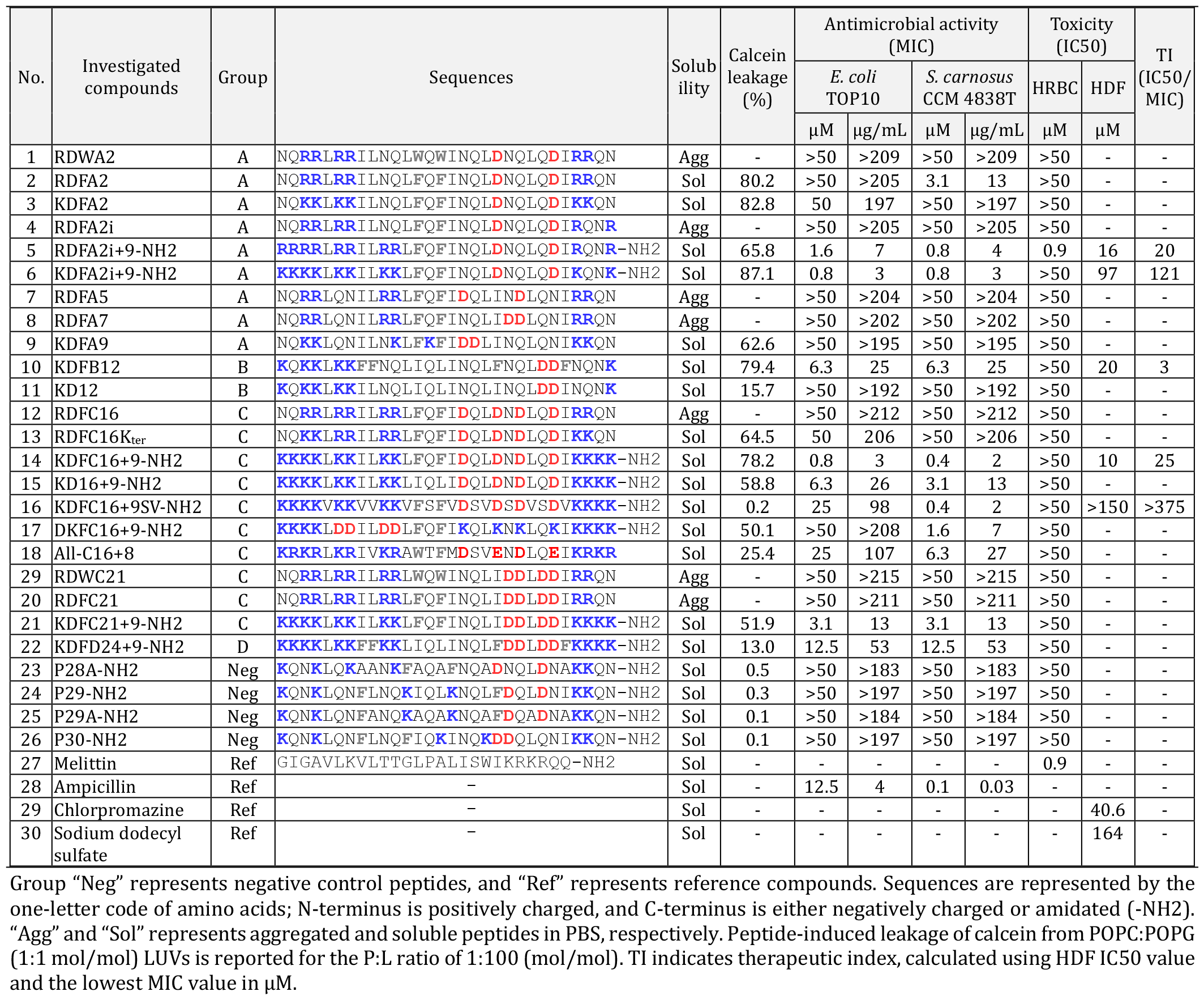
*In vitro* antimicrobial activity and human cell toxicity of *de novo* designed peptides.

Firstly, we performed the small molecule (calcein) leakage assay, which detects membrane permeabilization induced by peptides and is commonly used to indicate peptide pore formation (Figure 5a). Fourteen out of fifteen pore-stabilizing soluble peptides caused concentration-dependent leakage (efflux) of the fluorescent dye calcein (molecular weight 622.5 g/mol) from 0.02 mM POPC:POPG (1:1 mol/mol) large unilamellar vesicles (LUVs), used as a simple mimic of bacterial membranes. The only exception was the SV-variant peptide KDFC16+9SV-NH2, which did not cause dye leakage probably due to too narrow TBP formed with tightly packed luminal S residues^36^, observed in our simulation (Figure S2a-f). Leakage extent was >50% for 11 out of 14 peptides already at the lowest tested peptide/lipid (P:L) ratio of 1:100 (mol/mol), and the most active peptide, KDFA2i+9-NH2, caused ∼90% leakage (Table 1). The three peptides that caused <50% leakage were designed either: [1] with a lower amphipathicity due to a wide variety of amino acids (All_C16+9) or [2] with the narrowest pore cavity due to the strongest interactions at each peptide-peptide interface (KDFD24+9-NH2; compare with KDFC21+9-NH2) or [3] with not well stabilized TBP due to weaker peptide-peptide interactions with only 2 salt bridges at each interface and the absence of aromatic stacking (KD12; compare with KDFB12). These results suggest the chemical space covered by the design guidelines, show the trade-off between peptide-peptide interactions and pore cavity, and confirm that group A-B peptides require aromatic stacking for TBP stabilization. Furthermore, >50% leakage caused by KD16+9-NH2 confirmed that group C-D peptides form TBPs of optimal stability without aromatic stacking. Consistent with simulations, the leaky pore-forming ability was retained in designs with switched salt bridges (DKFC16+9-NH2) and in the presence of both inter- and intramolecular salt bridges (KDFA9). Note that the seven insoluble (pore-stabilizing) peptides had low net charge (+4 e) and R-clusters with a propensity for aggregation^41^ (Table 1). The insoluble peptides caused occasional leakage after partial dissolution via vigorous vortexing.

**Figure 5.**
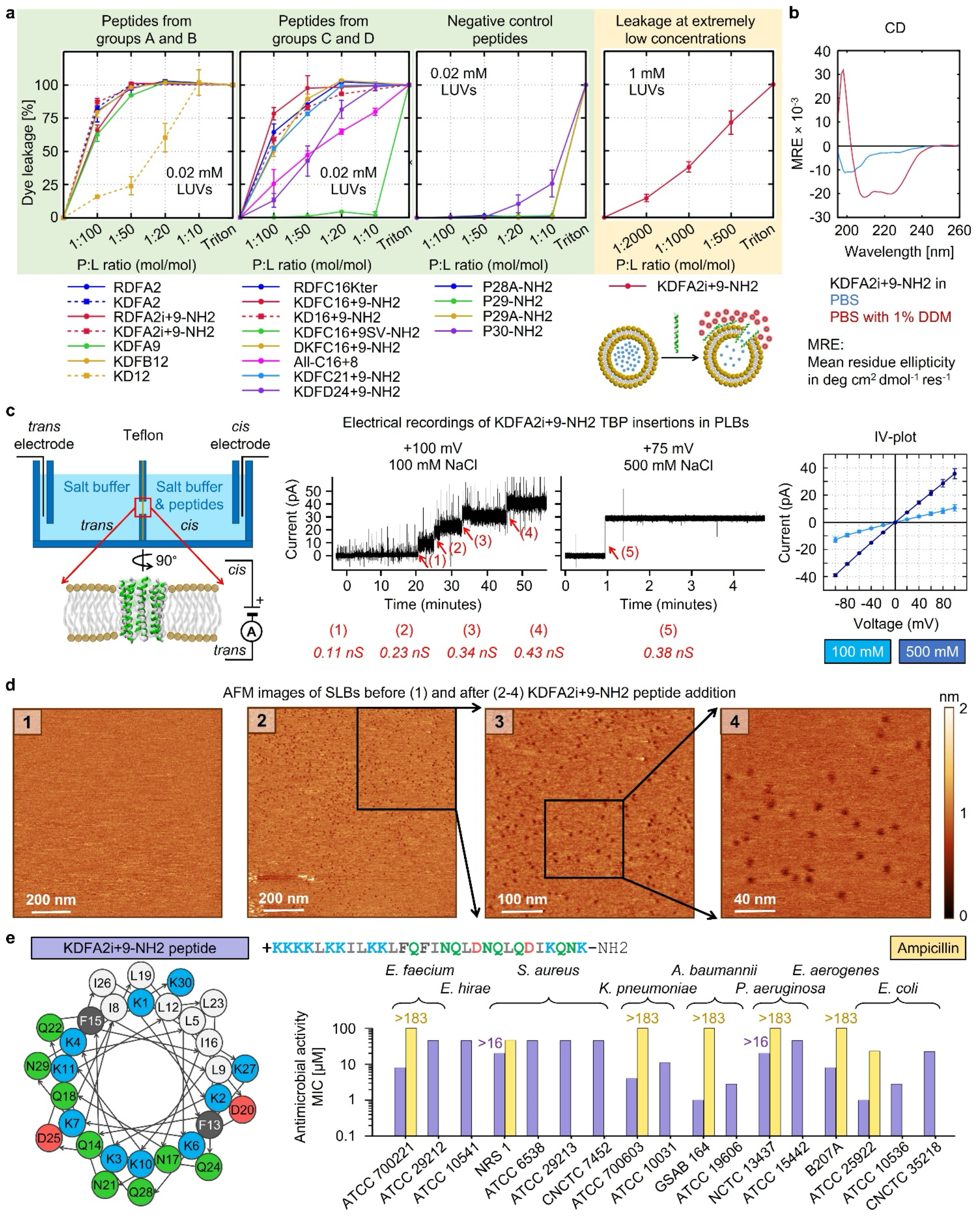
Pore-forming and antimicrobial activity of *de novo* designed peptides. (a) Concentration-depended peptide-induced leakage (efflux) of fluorescent dye calcein from POPC:POPG (1:1 mol/mol) LUVs. (b) CD spectra of peptides (20 μM) showing α-helical conformation in PBS with 1% DDM. (c: Left) Schematic illustration of electrophysiological assay. (c: middle) Current recordings showing insertions of KDFA2i+9-NH2 (500 nM) TBPs into DPhPC PLBs at two different salt concentrations. Insertion events are marked with red arrows and the conductance values are labelled. (c: Right) Current-voltage curves for singly inserted TBPs. (d) AFM topographic images showing membrane pore formation by KDFA2i+9-NH2 peptides (1 μM, P:L ratio 1:500 mol/mol) in POPC:POPG (1:1 mol/mol) SLBs. (e) Antimicrobial activity, reported as MIC, of KDFA2i+9-NH2 (helical wheel in left) and the reference antibiotic ampicillin against resistant ESKAPE bacteria.

Finally, as expected, the negative control peptides either did not permeabilize LUVs at all or in one case caused very low leakage (≤ 25%) at the highest tested P:L ratio of 1:10 (Figure 5a). Note that our simulations predicted that these peptides would not form stable TBPs, but the peptides could still form deformed or transient pores (Table S13) or may cause low leakage via local membrane thinning. Nevertheless, ∼25% leakage at P:L of 1:10 is significantly low compared to >80% leakage at P:L of 1:100 (10-fold lower concentration) caused by the template pore-forming peptides.

A barrel pore-forming peptide should be able to leak small molecules at a very low P:L ratio, as only a few nanopores are required to cause substantial leakage^33,42^. Indeed, the most active peptide, KDFA2i+9-NH2, caused noticeable leakage (∼20%) at an extremely low P:L of 1:2000, significant leakage (∼45%) at a very low P:L of 1:1000, and high leakage (∼75%) at a low P:L of 1:500 (Figure 5a). Note that leakage through few pores could be very slow and thus we measured leakage over a period of 1 hour after each peptide addition, close to the previous studies^33,42,43^. In addition, we used a higher lipid concentration of 1 mM to ensure that most of the peptides were bound to the LUVs^42,43^.

To verify that small molecule leakage was indeed caused by helical TBP formation, we performed channel electrical recordings using planar lipid bilayers (PLBs). We tested the most active peptide, KDFA2i+9-NH2, which, similar to other designed peptides, was α-helical in the model membranes (n-dodecyl β-D-maltoside (DDM) micelles and POPC:POPG LUVs), as confirmed by circular dichroism spectroscopy (Figures 5b and S6). Current recordings revealed stepwise pore-forming insertion events into DPhPC (1,2-diphytanolsn-glycero-3-phosphocholine) PLBs (Figures 5c and S7), where the peptides were pre-incubated with LUVs to form the oligomeric pore structures. Multiple pore insertions were observed over the 1 h study period, confirming formation of stable leaky TBP. Single-channel conductance varied linearly with the applied voltage, yielding a conductance of ∼0.1 nS and ∼0.4 nS at +100 mV in 100 mM and 500 mM NaCl, respectively. The conductance was then used to calculate pore radius using the equation^44^: 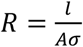. Considering a cylindrical nanopore, where *R* is the resistance (10^10^ S^-1^) with no surface charge, *l* is the membrane thickness (4 nm), *A* is the cross-section area (*πr*^*2*^), and *σ* is the bulk conductivity (1.06 Sm^-1^ in 100 mM NaCl^45^), we obtain a pore radius *r* of ∼4 Å, which is consistent with the radius of the MD simulated octameric TBP (Figure 3e).

To directly visualize KDFA2i+9-NH2 TBPs, we performed real-time AFM topographic imaging. At a low concentration of 1 μM (corresponding to a P:L ratio as low as 1:500 mol/mol), the peptides formed nanopores in POPC:POPG supported lipid bilayers (SLBs) (Figure 5d). The pores were detected within 20 minutes (imaging start time) of peptide addition and remained stable over the 3 hours study period. The number of pores gradually increased over time (Figure S8). Interestingly, the pore size appeared to be uniformly distributed around 6 nm in diameter, indicating structures larger than octamers, as predicted by our simulations of spontaneous pore formation (Figure S5). However, it should be noted that the estimated pore size is only approximate due to the size of AFM tip and membrane flexibility. At higher peptide concentrations (2-5 μM) we observed similar pore-like structures, including hexagonal lattice arrangements (Figure S9), similar to alamethicin pores^46^. We have also observed stacks of multiple bilayers demonstrating strong bilayer disruption. As expected, none of the 4 negative control peptides formed membrane pores (Figure S10), directly validating the leakage data.

To confirm the stabilization of larger TBPs as seen in AFM, we simulated TBPs of different sizes ranging from 4-mer to 16-mer and found that our designed interactions could indeed stabilize TBPs larger than octamer (Table S15 and Figure S11). To find the upper limit of the pore size, we further tested KDFA2i+9-NH2 TBPs by examining the transport of fluorescein isothiocyanate (FITC)-derivatized dextrans of different sizes across giant unilamellar vesicles (GUVs), an alternative membrane model for studying pore-forming proteins^20,47^. We observed the transport of 5 kDa (Stokes’ radius ∼1.7 nm) and 10 kDa (Stokes’ radius ∼2.3 nm) dextrans, but not 60-76 kDa (Stokes’ radius ∼5.4-6 nm), across the intact GUVs after peptide treatment, in contrast to the control GUVs without peptides (Figure S12). These results suggest that the designed TBPs could have a diameter between 5 and 10 nm.

### Antimicrobial application

To demonstrate potential antimicrobial applications of our de novo designed pore-forming peptides, we performed the broth microdilution assay using Gram-negative *Escherichia coli* TOP10 and Gram-positive *Staphylococcus carnosus* CCM 4838T (ATCC 51365), Biosafety Level 1 bacteria (Table 1). Out of 14 pore-forming soluble peptides, 12 showed antimicrobial activity against at least one of the two bacterial strains within 50 μM, the highest concentration tested. Two pore-forming peptides that showed no activity either had intramolecular salt bridges (KDFA9) or lacked aromatic stacking interactions (KD12), in addition to a low net charge (+4 e). Such behavior could be anticipated because electrostatic interactions, originating from peptide cationic character, were reported to play a crucial role in the initial attractive interactions of peptides with bacterial membranes^5,31,32^. In general, we found that increasing the net charge from +4 e to +9 e and/or R-to-K substitutions resulted in higher activity (compare R/KDFA2 and RDFC16Kter with R/KDFA2i+9-NH2 and KDFC16+9-NH2, respectively), in line with previous reports^5,31,32,35^. As expected, the 4 negative control peptides were inactive against both bacteria. Note that the list of 9 peptides that showed bactericidal activity against both bacterial strains includes the unusual SV-variant peptide, KDFC16+9SV-NH2 (Figure S13a). This suggests that the pores formed by KDFC16+9SV-NH2 were too narrow to allow calcein leakage, but wide enough to allow the leakage of bacterial cell material leading to cell death. Against *E. coli*, KDFA2i+9-NH2 and KDFC16+9-NH2 showed the highest activity with the minimum inhibitory concentration (MIC) of 0.8 μM, which was 15 times lower than the reference antibiotic ampicillin. Against *S. carnosus*, the most effective peptides were KDFC16+9-NH2 and KDFC16+9SV-NH2 with the MIC of 0.4 μM, which was 4 times higher than ampicillin.

### Toxicity

AMPs are required to be non-toxic against human cells for their effective pharmaceutical applications. Therefore, we tested all peptides for hemolysis toxicity against human red blood cells (HRBCs) (Table 1). Only one peptide, RDFA2i+9-NH2, showed hemolysis within 50 μM, the highest tested concentration (Figure S13b). RDFA2i+9-NH2 was the R-variant of the highly active AMP KDFA2i+9-NH2, and its half maximal inhibitory concentration (IC50) of 0.9 μM was similar to that of the reference hemolytic peptide melittin^48^. Other peptides remained non-hemolytic.

Next, we tested 5 peptides, RDFA2i+9-NH2, KDFA2i+9-NH2, KDFB12, KDFC16+9-NH2, and KDFC16+9SV-NH2, selected based on antimicrobial activity, for further toxicity against human dermal fibroblasts (HDFs) (Table 1). Peptides remained non-toxic at their antimicrobial concentrations and showed toxicity at concentrations at least an order of magnitude above the MIC values (Figure S13c). Note that peptide toxicity reduced after R-to-K substitutions (compare RDFA2i+9-NH2 with KDFA2i+9-NH2), probably due to weaker interactions with the zwitterionic phospholipids present in the outer leaflet of human plasma membranes^35^.

### Antimicrobial activity against ESKAPE pathogens

Our most active and yet non-toxic AMP, KDFA2i+9-NH2 (Table 1), was further tested against multidrug-resistant ESKAPE pathogens (Figure 5e), for which new antibiotics are needed^9^. Promising activity was obtained against *Acinetobacter baumannii* (GSAB 164), *E. coli* (ATCC 25922), and *Klebsiella pneumoniae* (ATCC 700603), with MICs 1, 1, and 4 μM, respectively. Susceptibility of these bacterial strains toward ampicillin was noticeably lower with MICs >183, 23, and >183, respectively. Peptide bactericidal activity was achieved already after 4 minutes (Table S16). The highest Therapeutic Index (TI) against ESKAPE pathogens was TI_(HRBCs-bacteria)_ >50 and TI_(HDFs-bacteria)_ ∼100, where a higher TI indicates higher bacterial selectivity over human.

Taken together, the results of AFM imaging, channel electrical recordings, small molecule leakage and macromolecule transport experiments are consistent with the results of MD simulations (CG and AA) and consistently support the peptide formation of stable, large, and functional TBPs. The formation of such leaky nanopores is therefore likely to be the mechanism of action of our designed AMPs.

## Discussion

Here, we employed a computer simulation-guided de novo design approach, evaluating ∼150 model peptides, to gain insight into the role of amino acids at each sequence position for the stabilization of octameric or larger TBPs. Single-ring TBPs smaller than octamer generally have pore diameters too small for the passage of molecules/solutes bigger than water and ions^15,16,18^. Octameric TBP was therefore the smallest and least computationally demanding system of choice, in line with the aim of our study. We used CG Martini 2.2 force field because of its computational efficiency and extensive use on similar systems. This model has its advantages and weaknesses thoroughly tested and well documented^49^ compared to the latest version, Martini 3, which still needs to be tested and improved in several aspects, e.g., peptide-peptide interactions^50^. The standard Martini 2.2 may overestimate protein-protein interactions^37^. We therefore combined the standard Martini with a scaled Martini to take advantage of their different characteristics. While the standard Martini identified the rough sequence patterns of peptides capable of stabilizing TBPs, the peptide-peptide interactions were then screened to identify the most stable interactions using the scaled Martini. Finally, we used the AA force fields to verify the results of the CG simulations. Using the computational approach, we have formulated a set of design guidelines that lead to de novo 52 sequence patterns with the capacity to generate ∼10^15^ peptides capable of forming TBPs.

Our designed TBPs have structural similarities with the pores/channels formed by natural peptide alamethicin^1^ and the other synthetically-derived peptides^2,14^, especially at the hydrophilic pore lumen allowing the passage of water, ions, and dye molecules, and the hydrophobic peptide patches interacting strongly not only with each other, but also with the membrane hydrophobic core, typical for a barrel-stave pore model^5,6^. There are several features which make our TBPs structurally stable and compact: (i) optimized salt bridge and aromatic stacking interactions, close-packing of residues and water-meditated hydrogen bonds, (ii) antiparallel peptide arrangements allowing pore-stabilizing attractive interactions between the oppositely charged neighboring transmembrane peptide termini, and hydrophilic peptide ends where cationic residues stabilize transmembrane orientation of peptides via attractive interactions with anionic lipid groups like phosphates.

Experimental determination of the high-resolution 3D structure of TBP is challenging due to the small size and dynamic nature of these structures. Therefore, we combined MD simulations with experiments to provide several lines of evidence for our designed TBP: (i) In CG MD, the designed peptides rearranged from a preformed membrane pore into the regular TBP that remained stable for 51 μs, additionally verified by 2 μs AA MD. In contrast, the negative control peptides, designed without the proposed intermolecular interactions, were unable to stabilize TBPs in both CG and AA MD and did not form pores in the fluorescence and AFM experiments. (ii) In spontaneous pore-forming MD, the designed peptides self-assembled into a regular TBP from the initial membrane surface-adsorbed state. (iii) AFM imaging allowed the direct visualization of the uniformly sized nanopores, which remained stable over the ∼3 hour study period, as expected for stable TBPs, since unstable pores are typically transient in nature and/or may vary in size as more peptides enter or leave the pore over the time^12,39,51,52^. (iv) Finally, upon addition of peptides pre-incubated in liposomes, channel conductance experiments showed multiple stepwise pore insertions over the 1 h study period, characteristic of stable and functional TBPs^14,16,18,20^.

Although our design was based on octameric TBPs, we noted that the formation of larger TBPs is also possible if the peptide-peptide interactions are able to adapt such arrangements^53^. Using the channel conductance data of ∼0.1 nS in 100 mM NaCl, we obtained a pore radius *r* of ∼4 Å, which is consistent with the radius of the MD simulated octameric TBP (Figure 3e). The pores observed in AFM appeared to be larger (diameter ∼6 nm) than the octamers and such large pores were also observed in spontaneous pore-forming MD. To determine the upper limit of the pore size, we performed the macromolecule (FITC dextran) transport assay, which suggested the formation of pores of sizes between 5 and 10 nm in diameter. However, it should be noted that the estimated pore size is highly approximate, as dextrans are flexible molecules that can deform and have a wide range of size distributions^47,54^. Taken together, the channel conductance, AFM imaging, small molecule leakage, and macromolecule transport experiments suggested that the designed peptides self-assemble into stable and functional TBPs with diameters of approximately 0.4-5 nm.

Our approach provides a molecular understanding of the relationship between sequence-specific interactions of α-helical peptides and the stability of assembled TBP structures with tunable channel. The identified role of individual residues in the proposed pore-forming sequence patterns provides control over pore properties and enables the custom design of peptide nanopores for different applications. Due to the increasing resistance of bacteria to conventional antibiotics, we chose to tune the designed TBP-forming peptides into selective antimicrobial compounds with rapid and direct killing action^6,8,9^. Peptide selectivity for bacteria over human cells can be increased by increasing the initial attractive electrostatic interactions^5,7^. Peptide net charge has been shown to be useful in this regard, as increased cationic character leads to more attractive interactions with negatively charged bacterial membranes than with the zwitterionic human membranes^31,32^. In the design, the hydrophilic peptide ends were made available for selectivity tuning. However, our simulations showed that charged residues should be added carefully in the ends of the pore-forming peptides, otherwise they may interfere with the pore-stabilizing interactions, leading to pore closure. Careful incorporation of additional cationic residues resulted in increased antibacterial activity, while maintaining the pore-forming activity and low toxicity unchanged, thus increasing selectivity. With that, designed peptides killed a broad spectrum of both Gram-negative and Gram-positive bacteria, including resistant ESKAPE pathogens, while causing negligible toxicity to human cells, offering a wide therapeutic window. The highest activity was obtained against the drug-resistant strains of *A. baumannii* and *E. coli* from the WHO priority pathogen list^9^. Therefore, our peptides can be a good starting point for the development of antibiotics.

Similar customization of the designed TBP-forming peptide sequences can be performed for other applications, e.g., anticancer agents^55^, single-molecule sensing and for sequencing of proteins and nucleic acids and in single-molecule chemistry for exploring sensitive chemical reactions^3,4^.

## AUTHOR INFORMATION

### Author Contributions

R.D. and R.V. conceived the project and designed research. R.D. carried out simulations and analyzed data. I.K. contributed with technical solutions to the problems in simulations. R.D. and R.V. designed the peptides and developed the design guidelines. R.D. conducted calcein leakage assay with help of I.K. R.D. and I.K. carried out broth microdilution and hemolysis assays. R.D. and J.P. performed AFM experiments. R.D. and E.V. performed channel electrical recordings. R.D. analyzed experimental data with help of I.K., J.P., E.V. and G.M. R.D. and R.V. wrote the manuscript with comments from all the authors.

### Notes

The authors declare the following competing financial interest(s): R.D., I.K., and R.V. are inventors on a EU Patent application filed by Masaryk University that covers the peptides and design guidelines described in this paper.

## ACKNOWLEDGMENT

We thank Martina Drabinová, Adelheid Hánačková, and Tereza Králová for help with experiments; Nada Labajová for advice on SLB formation; Guy Riddihough for help with the manuscript writing; Pavel Jungwirth for careful reading of the manuscript; and the members of Vácha group for discussions. This work was supported by the Czech Science Foundation (grant GA20-20152S), the European Research Council (ERC) under the European Union’s Horizon 2020 research and innovation programme (grant agreement No 101001470) and the project National Institute of virology and bacteriology (Programme EXCELES, ID Project No. LX22NPO5103) - Funded by the European Union - Next Generation EU. Additional support for experiments was provided by the project Development of the System of Commercialization of Results at Masaryk University TG02010067 financed by the Technology Agency of the Czech Republic. Computational resources were provided by the CESNET LM2015042 and the CERIT Scientific Cloud LM2015085 provided under the program Projects of Large Research, Development, and Innovations Infrastructures. For AFM experiments, we acknowledge CF Nanobiotechnology and Instruct-CZ Centre, supported by MEYS CR (LM2018127). For CD experiments, we acknowledge CF Biomolecular Interactions and Crystallography of CIISB, Instruct-CZ Centre, supported by MEYS CR (LM2023042) and European Regional Development Fund-Project “UP CIISB” (No. CZ.02.1.01/0.0/0.0/18_046/0015974). For fluorescence microscopy, we acknowledge CF CELLIM supported by MEYS CR (LM2023050 Czech-BioImaging).

